# Raf1 promotes successful Human Cytomegalovirus replication and is regulated by AMPK-mediated phosphorylation during infection

**DOI:** 10.1101/2023.07.26.550702

**Authors:** Diana M. Dunn, Ludia J. Pack, Joshua C. Munger

## Abstract

Raf1 is a key player in growth factor receptor signaling, which has been linked to multiple viral infections, including Human Cytomegalovirus (HCMV) infection. Although HCMV remains latent in most individuals, it can cause acute infection in immunocompromised populations such as transplant recipients, neonates, and cancer patients. Current treatments are suboptimal, highlighting the need for novel treatments. Multiple points in the growth factor signaling pathway are important for HCMV infection, but the relationship between HCMV and Raf1, a component of the mitogen-activated protein kinase (MAPK) cascade, is not well understood. The AMP-activated protein kinase (AMPK) is a known regulator of Raf1, and AMPK activity is both induced by infection and important for HCMV replication. Our data indicate that HCMV infection induces AMPK-specific changes in Raf1 phosphorylation, including increasing phosphorylation at Raf1-Ser621, a known AMPK phospho-site, which results in increased binding to the 14-3-3 scaffolding protein, an important aspect of Raf1 activation. Inhibition of Raf1, either pharmacologically or via shRNA or CRISPR-mediated targeting, inhibits viral replication and spread in both fibroblasts and epithelial cells. Collectively, our data indicate that HCMV infection and AMPK activation modulate Raf1 activity, which are important for viral replication.

**Importance:** Growth factor signaling plays a critical role in many aspects of viral infection. Here we show that a component of one of these pathways, Raf1, contributes to successful infection of Human Cytomegalovirus (HCMV). We find that AMP-activated protein kinase (AMPK), which is known to be important for HCMV infection, modulates Raf1 phosphorylation throughout infection, and contributes to Raf1 binding to its activating co-factor, 14-3-3. In addition, inhibition of Raf1 inhibits HCMV infection and viral spread. These results suggest a link between two cellular pathways that are important for HCMV replication, AMPK signaling and growth factor receptor signaling, that converge as an important aspect of HCMV infection. This could lead to the potential for new therapeutic targets in immunocompromised individuals afflicted by acute HCMV infection.

## Introduction

Human Cytomegalovirus (HCMV) infection is a prevalent virus infecting approximately 60-90% of the global population (1). Although it remains latent in healthy individuals, it can lead to damaging acute infection in immunocompromised individuals such as neonates (2, 3), transplant recipients (4, 5), those undergoing immunosuppressive anti-cancer therapies (6–10), and those receiving anti-AIDS therapies (11). Current treatments for HCMV are suboptimal, often resulting in drug resistance and unwanted side effects (12–15). There is therefore a need for the development of novel therapeutics to treat acute HCMV infection.

HCMV is a beta herpes virus with a large double-stranded DNA genome encoding for upwards of 200 open reading frames with the potential to express hundreds of viral proteins, many of which have unknown functions or expression patterns (16). HCMV, like many other viruses, hijacks and relies on various host factors for successful infection (17). One of these host proteins is AMP-activated protein kinase (AMPK), a central metabolic stress regulator, whose expression and activity are induced by HCMV infection to support virally-induced metabolic remodeling and promote viral replication (18, 19).

Growth factor signaling has been linked to the productive infection of multiple viruses including influenza (20), Zika (21), coronaviruses (22), and cytomegalovirus (23) Growth factor receptors facilitate viral entry, internalization, replication, and egress (24) A typical growth factor signaling pathway involves binding to receptor tyrosine kinases such as platelet-derived growth factor receptor (PDGFR) or epidermal growth factor receptor (EGFR), which results in downstream activation of Ras and the subsequent cascade of Raf1, MEK, and ERK leading to regulation of multiple cellular processes including proliferation, survival, differentiation, migration, apoptosis and more (25, 26) (**Figure 1A**).

**Figure 1.**
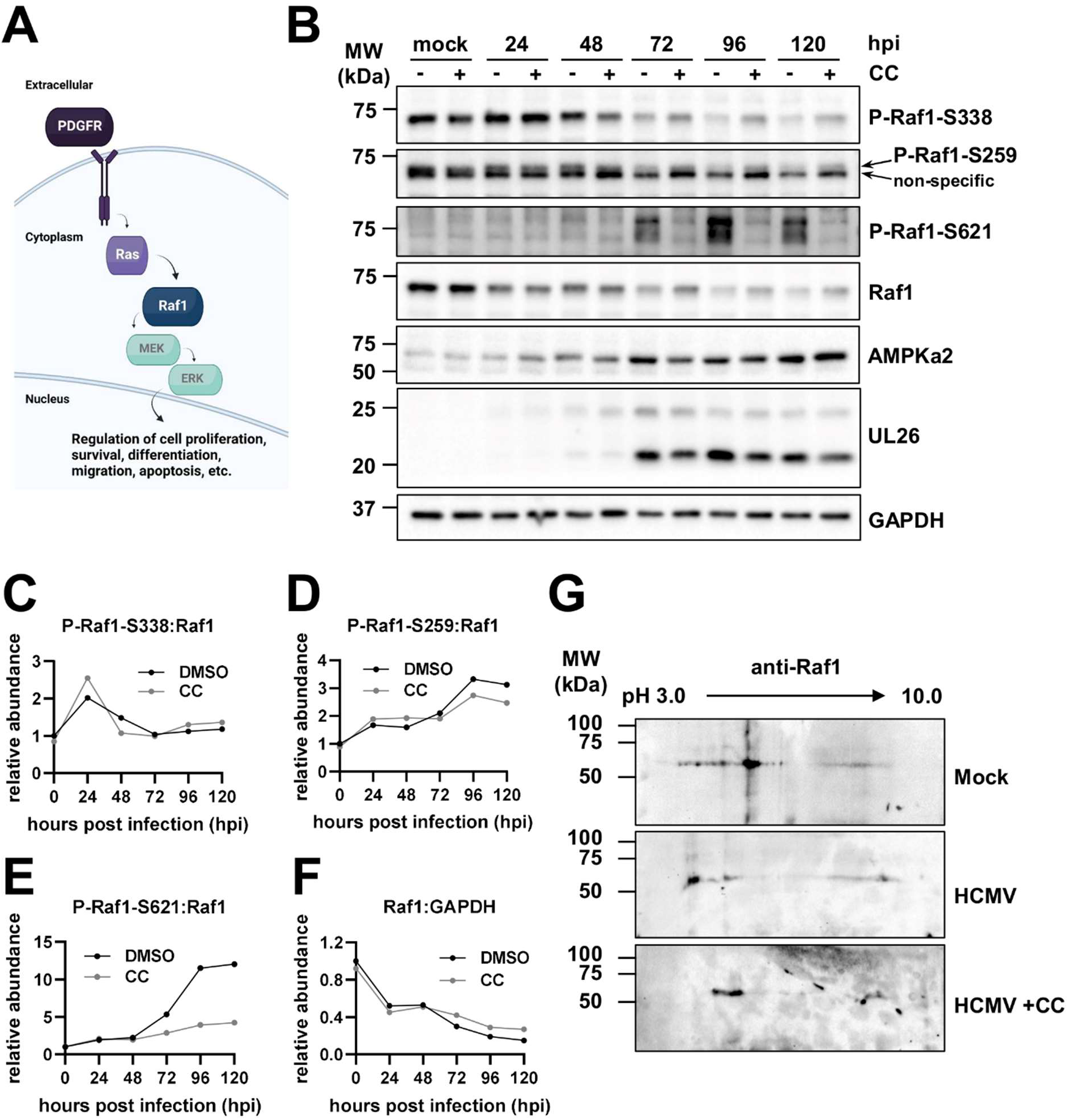
HCMV infection induces AMPK specific phosphorylation of Raf1. (A) Schematic depicting the Raf1 signaling pathway. (B) Western blot analysis of Raf1 over the course of infection (120 hours) in the absence and presence of Compound C (CC) in MRC5 fibroblasts. GAPDH was used as a loading control and UL26 is a control for successful viral infection. (C-F) Quantification of Western blots in (B), where zero hours post-infection (hpi) is representative of mock infection, are comparing phospho-Raf1 to total Raf1 for (C) P-Raf1-S338, (D) P-Raf1-S259 (top band), and (E) P-Raf1-S621, and (F) total Raf1 levels normalized to GAPDH. (G) Two-dimensional (2-D) gel electrophoresis of Raf1 during mock infection and HCMV infection with or without CC treatment, MOI = 3.0 at 48 hpi in MRC5 fibroblasts.

Human Cytomegalovirus (HCMV) relies on multiple stages of growth factor signaling to support productive infection (23). PDGFR and EGFR not only facilitate viral entry (27, 28), but also signal downstream cellular pathways to sustain viral signaling, trafficking, gene transcription, and latency (29–31). Soluble derivatives of PDGFR have been demonstrated to bind and neutralize free HCMV particles, thus reducing viral entry into cells (32, 33). Further, inhibition of PDGFR by imatinib or nilotinib can inhibit viral entry and HCMV infection when cells are pretreated with these drugs (34). Likewise, treatment of cells with other various broad-spectrum MAPK inhibitors attenuates HCMV replication (35, 36). However, it is well known that HCMV is unable to replicate in transformed cell lines, and Ras/SV40-mediated transformation of human cells leads tothe inability of HCMV to replicate (37). Collectively, these data indicate a complex relationship between HCMV and growth factor receptor signaling.

Raf1, also known as c-Raf, is a heavily phosphorylated Mitogen-Activated Protein Kinase Kinase Kinase (MAPKKK), and the dynamic nature of its phosphorylation plays a complicated role in its kinase activity (38). Phosphorylation at Ser338 activates Raf1 (39), while phosphorylation at Ser259 results in Raf1 inhibition (40). Raf1 Ser621 phosphorylation is necessary for its activation and stability, achieved through binding to the scaffold protein and activating co-factor, 14-3-3 (41–45). The main site within Raf1 that is targeted by AMPK is Ser621, although Ser259 is also putatively modified by AMPK as well as other kinases, including PKCA and Raf1 itself (40, 46, 47). Crosstalk between these phosphorylation events and others within Raf1 can impact multiple aspects of the protein including its cellular location, stability, and activity (38, 44, 48).

Here, we explore the phosphorylation of Raf1 and its contributions to HCMV infection. We find that HCMV induces AMPK-specific phosphorylation of Raf1 and in turn, increases Raf1 binding to the scaffolding protein 14-3-3. Pharmaceutical inhibition of Raf1 results in reduced viral DNA and protein accumulation and attenuates infectious virion production. Further, Raf1 knockdown by shRNA or knockout by CRISPR results in decreased viral titers and reduced cell-to-cell spread.

## Results

### HCMV infection induces AMPK-specific phosphorylation of Raf1

To evaluate the effects of AMPK on Raf1 we first assessed the specific phosphorylation of Raf1-S338, -S259, and -S621 by western blot during infection in the presence of DMSO or AMPK inhibitor Compound C (CC) (**Figure 1B**). There was an early increase in activated Raf1 phosphorylated at S338, which dropped as infection proceeded (**Figure 1B-C**). The ratio of Raf1-S338 phosphorylation to total remained unchanged in DMSO versus Compound C-treated samples at each time point, suggesting that AMPK does not impact the phosphorylation of Raf1 at this site (**Figure 1C**). Although difficult to quantify due to the quality of available antibodies, the ratio of Raf1-S259 phosphorylation increased during infection but decreased modestly upon Compound C treatment (**Figure 1D**). In contrast, the level of Raf1-S621 phosphorylation increased substantially during infection, which was blocked by treatment with Compound C (**Figure 1E**). Notably, Raf1 levels dropped by approximately 50% relative to mock at 24 hours post-infection (hpi) and remained there until 48 hpi, then decreased further as infection progressed (**Figure 1F**). Although total Raf1 levels were marginally higher with Compound C treatment compared to DMSO control at each time point, they were not fully restored to mock or early infection levels (**Figure 1F**). The increase in Raf1-S621 phosphorylation and loss of total Raf1 expression mirrored increases in AMPKa2 protein levels, which have previously been found to be important for HCMV infection (**Figure 1B**) (18, 19).

To further explore how HCMV and AMPK impacted the post-translational modifications of Raf1, we analyzed Raf1 via two-dimensional (2-D) gel electrophoresis (**Figure 1G**). During HCMV infection, Raf1 populations had a significantly higher negative charge relative to Raf1 during mock infection, resulting in a shift toward pH 3.0, consistent with increased phosphorylation (**Figure 1G**). The addition of Compound C during infection resulted in less negatively charged Raf1, shifting the Raf1 isoforms towards the more basic (pH 10.0) cathode, suggesting substantially decreased phosphorylation (**Figure 1G**). These results indicate that AMPK likely plays an important role in regulating Raf1 phosphorylation during HCMV infection.

Given the observation that HCMV induced increases in Raf1-S621 phosphorylation (**Figure 1**), we wanted to explore the impact of the expression of a Raf1 allele that is non-phosphorylatable at this site on HCMV replication. Towards this end, Flag-Raf1-WT or Flag-Raf1-S621A were constructed and delivered to fibroblasts via lentiviral transduction (**Figure 2**). Raf1 protein levels were elevated compared to empty vector control cells (EV) (**Figure 2A**). In addition, Flag-Raf1-WT accumulated to higher levels than Flag-Raf1-S621A, which could be expected as S621 phosphorylation is important for Raf1 stability (43). Overexpression of either Flag-Raf1-WT or Flag-Raf1-S621A also increased S338 phosphorylation (**Figure 2A**), suggesting overexpression of Raf1 induces its activity in our fibroblasts. Expression of P-ERK was also elevated, indicative of an active Raf1 pathway. Phosphorylation of S621 was elevated in the WT-Raf1 cell line, but not in the cells expressing Raf-S621A confirming that this mutation attenuates phosphorylation at this site (**Figure 2A**).

**Figure 2.**
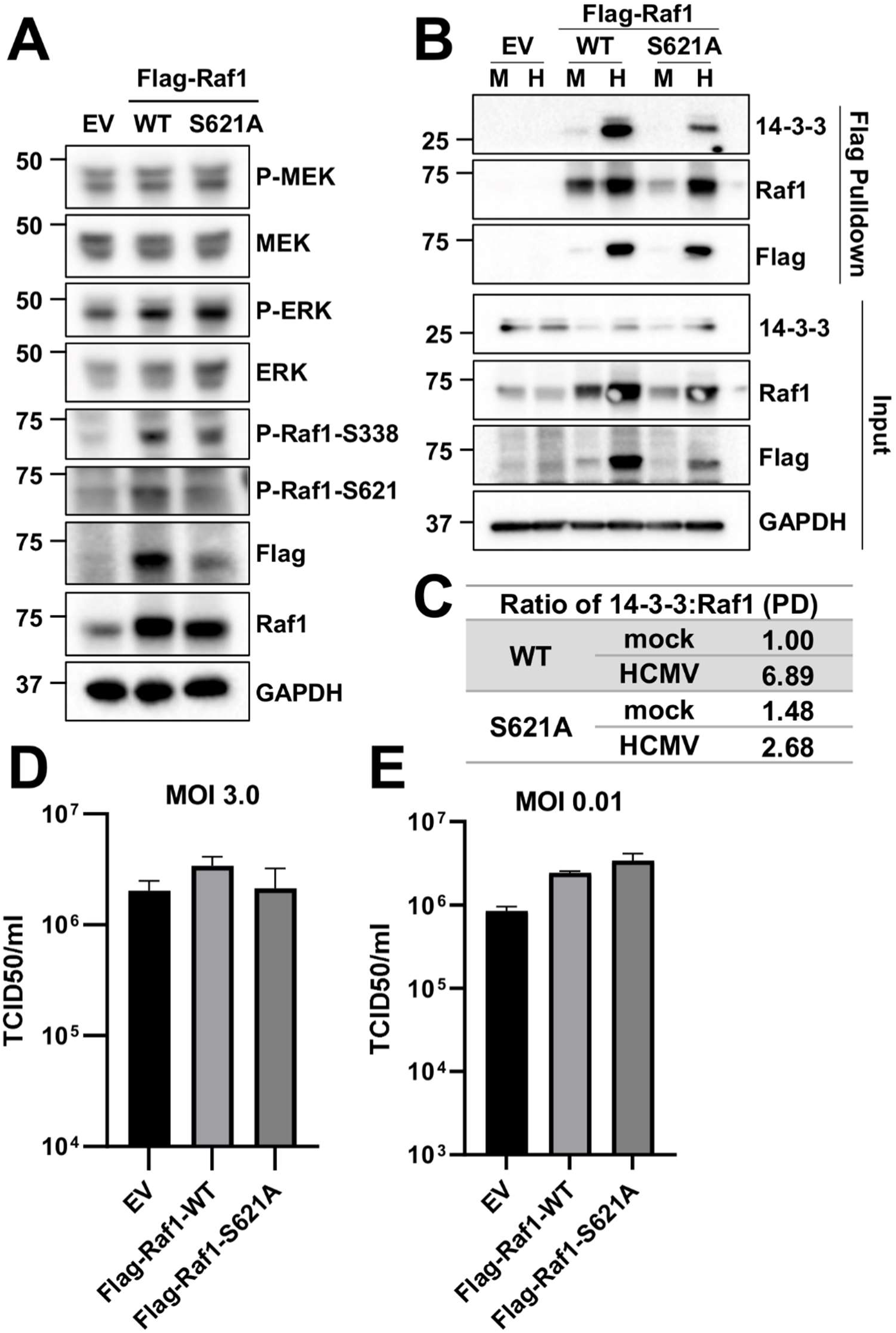
Phosphorylation of S621 enhances Raf1 binding to 14-3-3 during HCMV infection. (A) Overexpression of pLenti-Flag-Raf1-WT or pLenti-Flag-Raf1-S621A were compared to pLenti-empty vector (EV) control MRC5 fibroblasts and assessed by Western blot using antibodies specific to flag, Raf1 and S621- or S338-phosphorylated Raf1, and total or phospho-MEK and -ERK. GAPDH was used as a loading control. (B) EV, Flag-Raf1-WT and -S621A cells were mock (M) or HCMV (H) infected with AD169-WT at an MOI of 3.0 for 72 hours prior to Flag-pulldown and co-precipitation of endogenous 14-3-3. (C) Ratio of 14-3-3 to Raf1 in (B) was quantified. (D-E) Cells were infected at an MOI of 3.0, for 96 hours (avg±SEM, n=3) (D), or an MOI of 0.01 for 10 days (avg±SEM, n=6) (E), and viral titers were assessed by TCID50.

Raf1-S621 phosphorylation is a prerequisite for its activation via binding to the scaffold protein 14-3-3 (41–45). We, therefore, assessed the interaction between Raf1 and 14-3-3. Cells were transduced with either EV, Flag-Raf1-WT, or Flag-Raf1-S621A expression constructs, and subsequently infected (MOI 3.0). At 72 hours post-infection, the Flag-fused Raf1 proteins were purified via Flag-antibody affinity and analyzed for co-precipitation with 14-3-3 (**Figure 2B**). There was a significant increase in the WT-Raf1 association with 14-3-3 during HCMV infection (**Figure 2B-C**). This association was significantly reduced when the Raf1-S621A allele was expressed (**Figure 2B-C**), consistent with previous reports indicating S621 phosphorylation as being important for 14-3-3 binding (41–45). These data suggest that AMPK-mediated phosphorylation of Raf1 increases its binding to 14-3-3 during HCMV infection, thus regulating Raf1 stabilization and activation.

Next, we assessed how the expression of WT-Raf1 or Raf1 containing the S621A mutation impacted HCMV infection. At a high MOI (3.0), neither expression of WT nor the mutant Raf1 allele significantly impacted viral titers (**Figure 2D**). Similarly, at a low MOI (0.01), there was no substantial impact on the production of HCMV progeny after expressing either Raf1 allele (**Figure 2E**). These data suggest that expression of either WT or S621A-mutated Raf1 does not impact HCMV infection. However, firm conclusions about the role of AMPK-mediated Raf1 phosphorylation at S621 remain difficult given the presence of endogenous Raf1, which presumably remains capable of S621 phosphorylation.

### Pharmacological inhibition of Raf1 reduces HCMV infection

We next evaluated the effect of Raf1 inhibition on viral replication. Regorafenib and Sorafenib are Raf1-specific inhibitors, although they also inhibit other kinases in the Raf1 pathway, including b-Raf, VEGFR1, VEGFR2, VEGR-3, PDGFRB, Flt-3 and cKIT, albeit with higher IC50s (**Table 1**) (49, 50). To examine how HCMV replicated in the presence of these compounds we monitored HCMV infection in two fibroblast cell lines, MRC5 and HFF, in response to titrated compounds (**Figure 3**). After 5 days, cells positive for GFP were counted and plotted per area of cell growth against compound concentration (**Figure 3A-D**). Both Sorafenib and Regorafenib limited viral replication, as indicated by the loss of GFP-positive cells at 2.17 uM (**Figure 3A-D**). These results were consistent with a previous report using Sorafenib (36). We subsequently examined how these compounds impact the production of viral progeny and found that the amount of infectious virus produced in their presence was below the limit of detection (**Figure 3E-F**). As expected, treatment with either Sorafenib or Regorafenib substantially reduced the phosphorylation of downstream Raf1 targets, including p-MEK and p-ERK (**Figure 3G-H**). Further, the inhibitors were generally well-tolerated in mock and HCMV-infected cells as indicated by consistent nuclear counts (**Figure 4A-D**). Together, these data indicate inhibition of the Raf1 pathway can inhibit viral infection and spread without impacting cell viability.

**Table 1.** Reported IC50s of Regorafenib and Sorafenib to Raf1 pathway proteins {Wilhelm, 2004 #2778;Wilhelm, 2011 #2779}.

| Raf family protein | IC50 – Regorafenib | IC50 – Sorafenib |
| --- | --- | --- |
| Raf1 | 2.5 nM | 6 nM |
| B-Raf | 28 nM | 22 nM |
| B-Raf (V600E) | 19 nM | 38 nM |
| PDGFRB | 22 nM | 57 nM |

**Figure 3.**
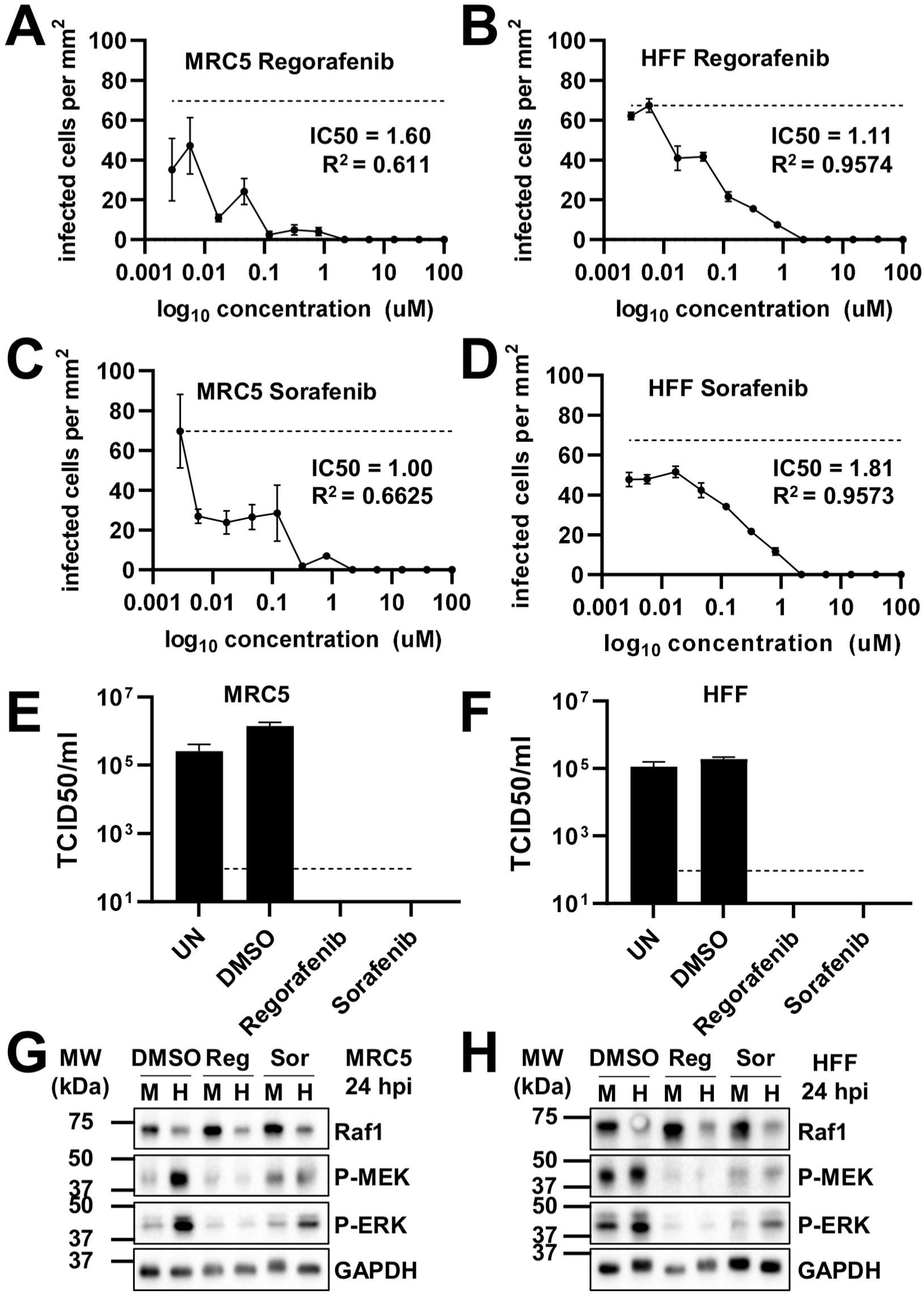
Pharmacological inhibition of the Raf1 pathway inhibits HCMV infection. (A-B) Regorafenib (Reg) or (C-D) Sorafenib (Sor) treated (A,C) MRC5 or (B,D) HFF fibroblasts were treated with indicated concentration of drugs or DMSO control at the time of infection. (A-D) Cells were infected at an MOI of 0.1 with AD169-GFP for 5 days, then fixed. Using the Cytation and Gen5 software, infected GFP positive cells were counted per area of the well and plotted against drug concentration (avg±SEM, n=3). IC50 values were calculated in GraphPad Prism. Dotted line indicates average maximal GFP in untreated cells. (E-H) Fibroblasts were infected at an MOI of 3.0 in the presence of DMSO or 2.17 uM Regorafenib or Sorafenib, added at the time of viral adsorption. UN is an untreated control. Viral titers from (E) MRC5 or (F) HFF cells, were assessed by TCID50 (avg±SEM, n=3), 120 hpi, the dotted line indicates the limit of detection. In (G) MRC5 or (H) HFF cells, at 24 hours post-infection (hpi), phosphorylation of MEK and ERK was assessed by western blot.

**Figure 4.**
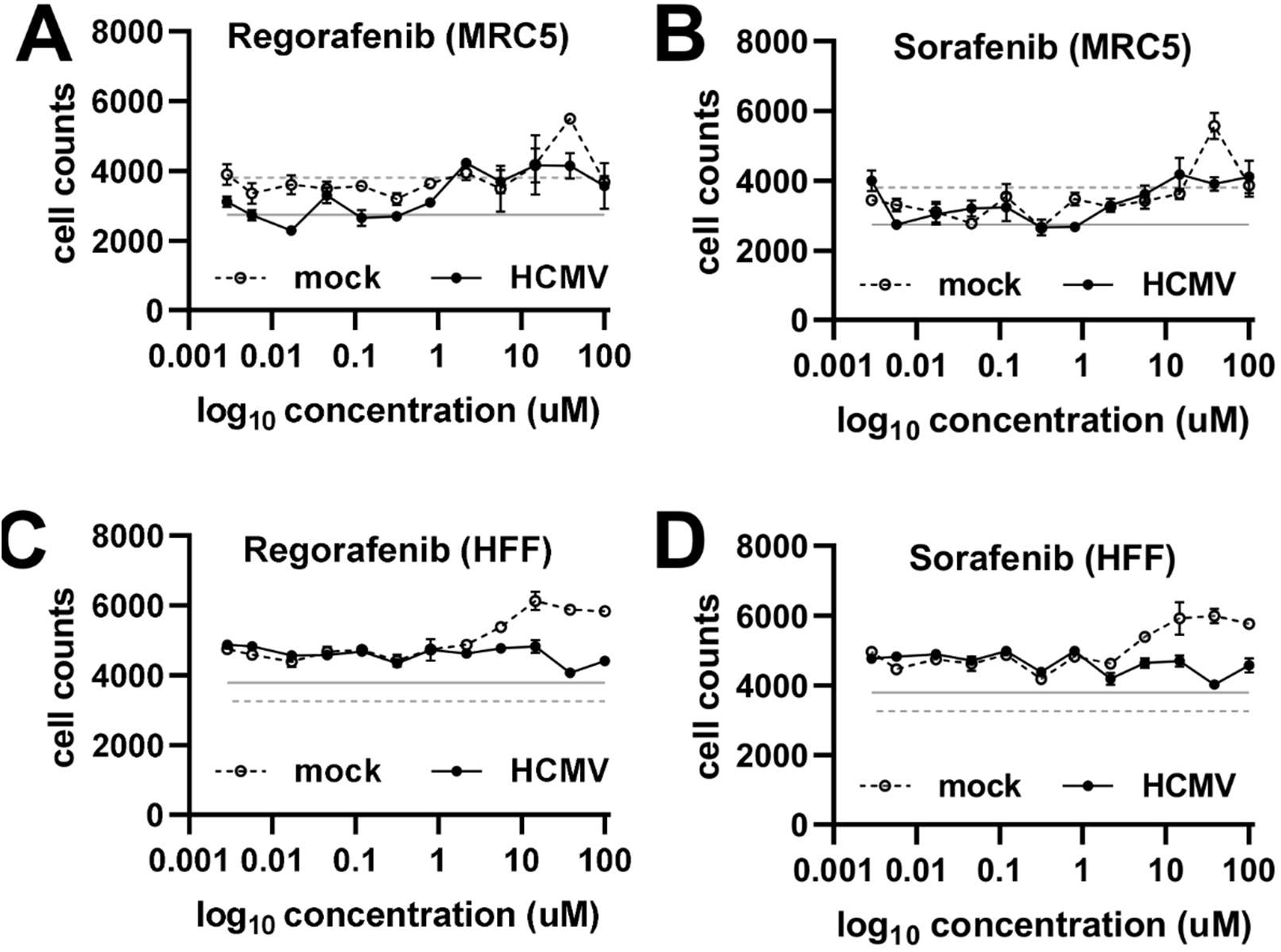
Pharmacological inhibition of the Raf1 pathway does not impact fibroblast cell counts. (A-B) MRC5 or (C-D) HFF fibroblasts were treated with indicated concentration of (A, C) Regorafenib or (B, D) Sorafenib at time of infection, AD169-GFP, MOI = 0.1, fixed on Day 5. (A-D) Using the Cytation and Gen5 software, Hoechst-stained nuclei were counted and plotted against concentration (avg±SEM, n=3). Solid gray line indicates average number of nuclei in untreated HCMV infected cells and dotted gray line indicates average number of nuclei in untreated mock infected cells.

### Inhibition of Raf1 negatively impacts viral DNA and protein accumulation

To elucidate how Raf1 inhibition impacted the viral life cycle, we first measured viral DNA accumulation from 48 to 120 hours post-infection (hpi) (**Figure 5A**). Significantly less viral DNA accumulated over the course of infection when cells were treated with Regorafenib and Sorafenib relative to DMSO controls, as indicated by the area under the curve for each sample (**Figure 5B**). Regorafenib and Sorafenib treatment modestly reduced IE1, IE2, and UL44 protein levels at 24 hpi and 72 hpi, but by 96 hpi, inhibitor treatment resulted in substantially less accumulation of these proteins as well as pp28, a late protein (**Figure 5C**). The modest impact of Raf1 inhibition on immediate early gene expression suggests that Raf1 activity is not necessary for viral entry, but rather indicates that Raf1 is important for post-entry early events of infection leading to DNA replication.

**Figure 5.**
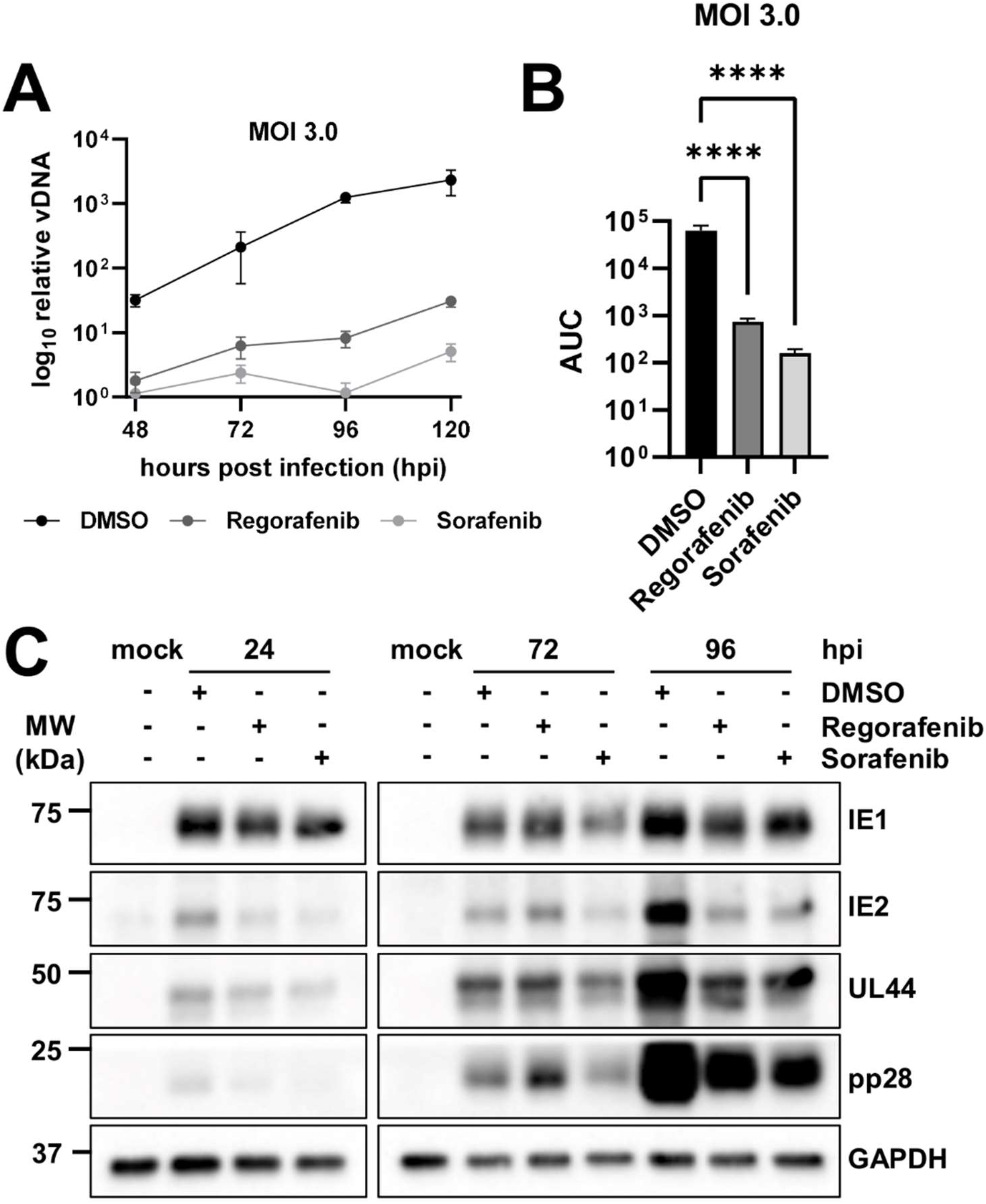
Inhibition of the Raf1 pathway negatively impacts viral DNA and protein accumulation. (A-C) MRC5 fibroblasts were treated with 2.17 uM of Regorafenib or Sorafenib, or DMSO control at the time of infection with AD169, at an MOI of 3.0 for indicated amount of time. (A) Viral DNA was quantified by qPCR of the viral gene, IE1 (avg±SD, n=12). (B) Samples in (A) were compared by area under the curve (AUC) to show the significance between treatments over the course of infection (avg±SD, n=12). (C) Viral protein expression levels at 24-, 72-, and 96 hours post-infection (hpi) were assessed by western blot. GAPDH was used as a loading control.

### Knockdown or partial knockout of Raf1 reduces HCMV infection

Given that Sorafenib and Regorafenib can both inhibit other kinases in the Raf1 pathway (**Table 1**), we employed other methods to analyze Raf1’s potential contributions to HCMV infection. First, we targeted Raf1 with two different shRNA constructs in HFF cells. The knockdown efficiency was assessed by qPCR, which indicated a 54% and 58% loss in Raf1 RNA expression with constructs R1-1 and R1-5 respectively, compared to the empty vector (EV) shRNA control (**Figure 6A**). At a high MOI (3.0), these constructs reduced HCMV infection by approximately 10-fold (**Figure 6B**). A similar replication defect was observed at a low MOI (0.01) (**Figure 6C**), highlighting the importance of Raf1 for HCMV infection.

**Figure 6.**
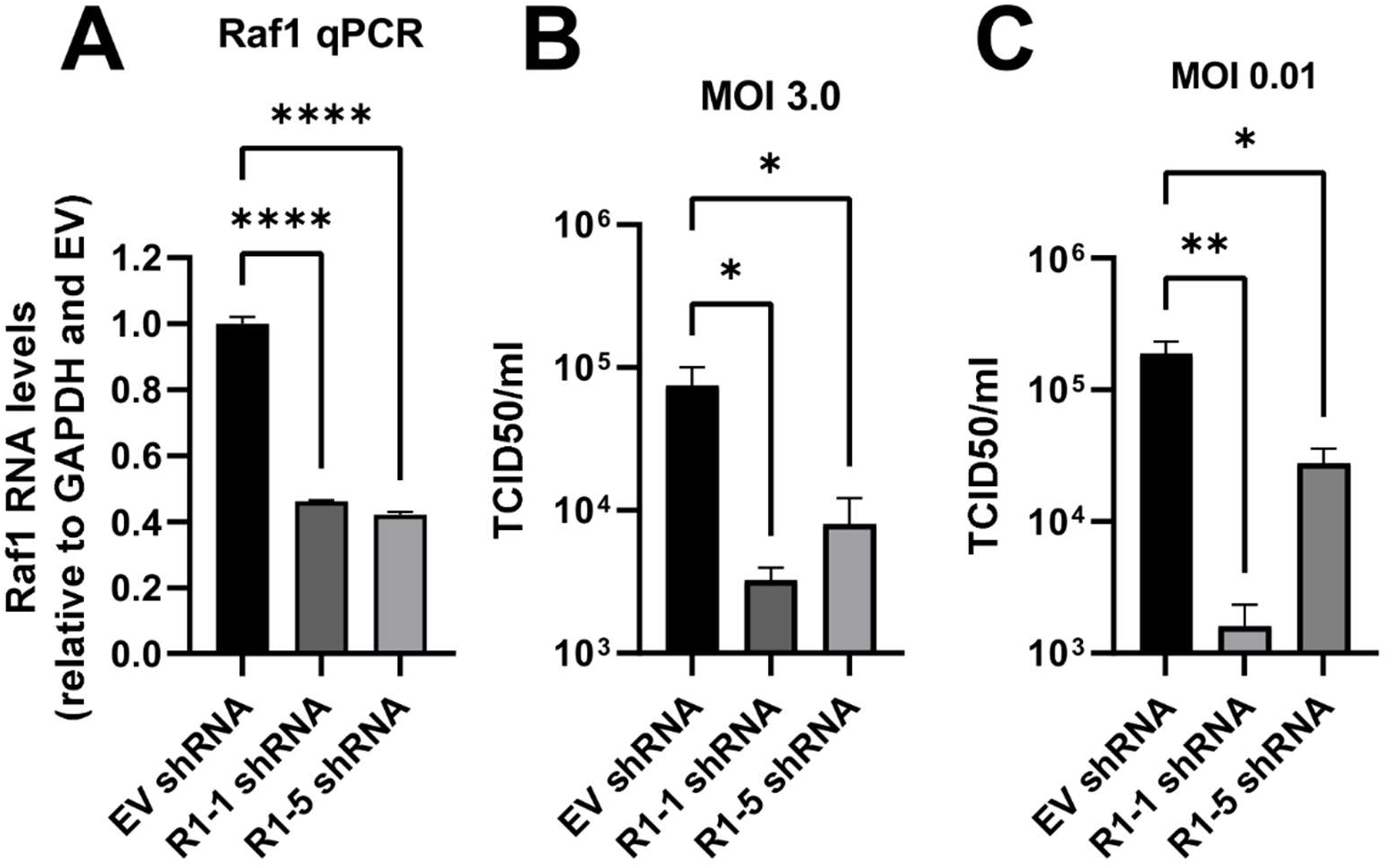
Raf1 shRNA-mediated knockdown inhibits HCMV infection. (A-C) Raf1 was knocked-down via lentiviral delivery of shRNA guides (R-1-1 and R1-5). (A) Knockdown efficiency in uninfected HFF fibroblasts was assessed by qPCR (avg±SEM, n=4), RNA levels were normalized to GAPDH and empty vector (EV) control. (B-C) Virus was harvested, and titer assessed by TCID50 (avg±SEM, n=4) at a (B) high MOI of 3.0 at 120 hours post-infection, and (C) a low MOI of 0.01 at 10 days post infection.

Additionally, we generated polyclonal Raf1 knockout (KO) cell lines via transfection of ribonucleoprotein-CAS9 and Raf1-specific guide RNAs (sgRNA) into MRC5 and HFF cells. The Raf1 KO score was 83% in the MRC5 cell line and 53% and 85% in the HFF #1 and #2 cell lines respectively (**Figure 7A**). Consistent with these findings, CRISPR targeting of Raf1 resulted in substantially reduced Raf1 levels in the MRC5 KO cells (**Figure 7B**). During infection, CRISPR-mediated inactivation of Raf1 in the MRC5 cells did not impact the accumulation of IE1 or UL26 (**Figure 7B**). However, we observed that Raf1 KO in MRC5 cells reduced viral titers relative to parental cells at 120 hours post-infection (**Figure 7C**). We also assessed how Raf1 knockout impacted HCMV cell-to-cell spread in both Raf1 KO MRC5 and HFF cells. After infection at an MOI of 0.05, and incubation for 10 days, the number of GFP-positive cells was significantly lower in the Raf1 KO cells (**Figure 7D-E**). Similar to the treatment with pharmacological inhibitors and Raf1-shRNA, these results indicate that Raf1 is important for high-titer HCMV replication.

**Figure 7.**
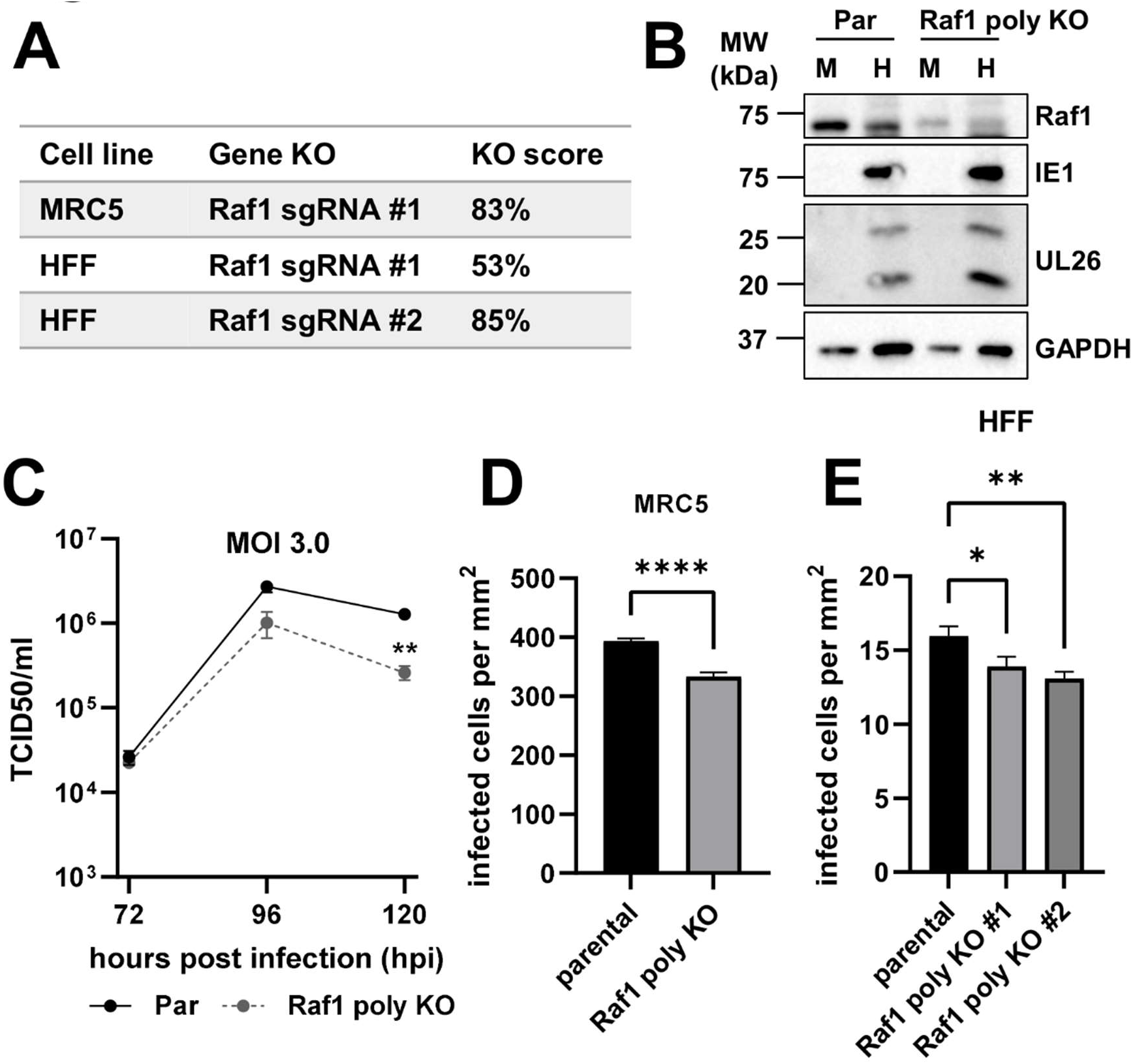
CRISPR-mediated partial knockout of Raf1 attenuates cell to cell spread of HCMV. (A) Knockout (KO) scores for CRISPR Raf1 KO cell lines generated. (B-D) MRC5 parental (Par) and Raf1 polyclonal CRISPR knockout (poly KO) cells were infected at an MOI of 3.0 for (B) 96 hours and protein expression assessed by western blot during mock (M) or HCMV (H) infection. (C) At 72-, 96-, and 120 hours post-infection (hpi), viral titers were assessed by TCID50 (avg±SEM, n=2). Significance is based on a student’s t-test at 120 hpi. (D) MRC5 (avg±SEM, n=72) and (E) HFF (avg±SEM, n=48), parental and Raf1 polyclonal CRISPR knockout cell lines were infected with AD169-GFP at an MOI of 0.05 for 10 days. Infected cells per area of the well were counted via GFP expression.

### Pharmacological inhibition of Raf1 attenuates HCMV TB40/E cell-to-cell spread in fibroblasts and epithelial cells

So far, our experiments have employed the use of a laboratory strain of HCMV, AD169. To further test the impact of Raf1 inhibition on the replication of a more clinically relevant HCMV strain, we targeted Raf1 during TB40/E infection in fibroblasts and epithelial cells. First, viral spread was measured using the AD169-GFP virus in MRC5 fibroblasts, where inhibition of Raf1 by Regorafenib and Sorafenib significantly inhibited viral cell-to-cell spread (**Figure 8A**). In the same cells, the cell-to-cell spread of TB40/E was also significantly reduced (**Figure 8B**). Similarly, the cell-to-cell spread of TB40/E was reduced in ARPE19 epithelial cells treated with Raf1 inhibitors relative to DMSO (**Figure 8C**). These data suggest that Raf1 is important for multiple strains of HCMV to replicate in both fibroblasts and epithelial cells.

**Figure 8.**
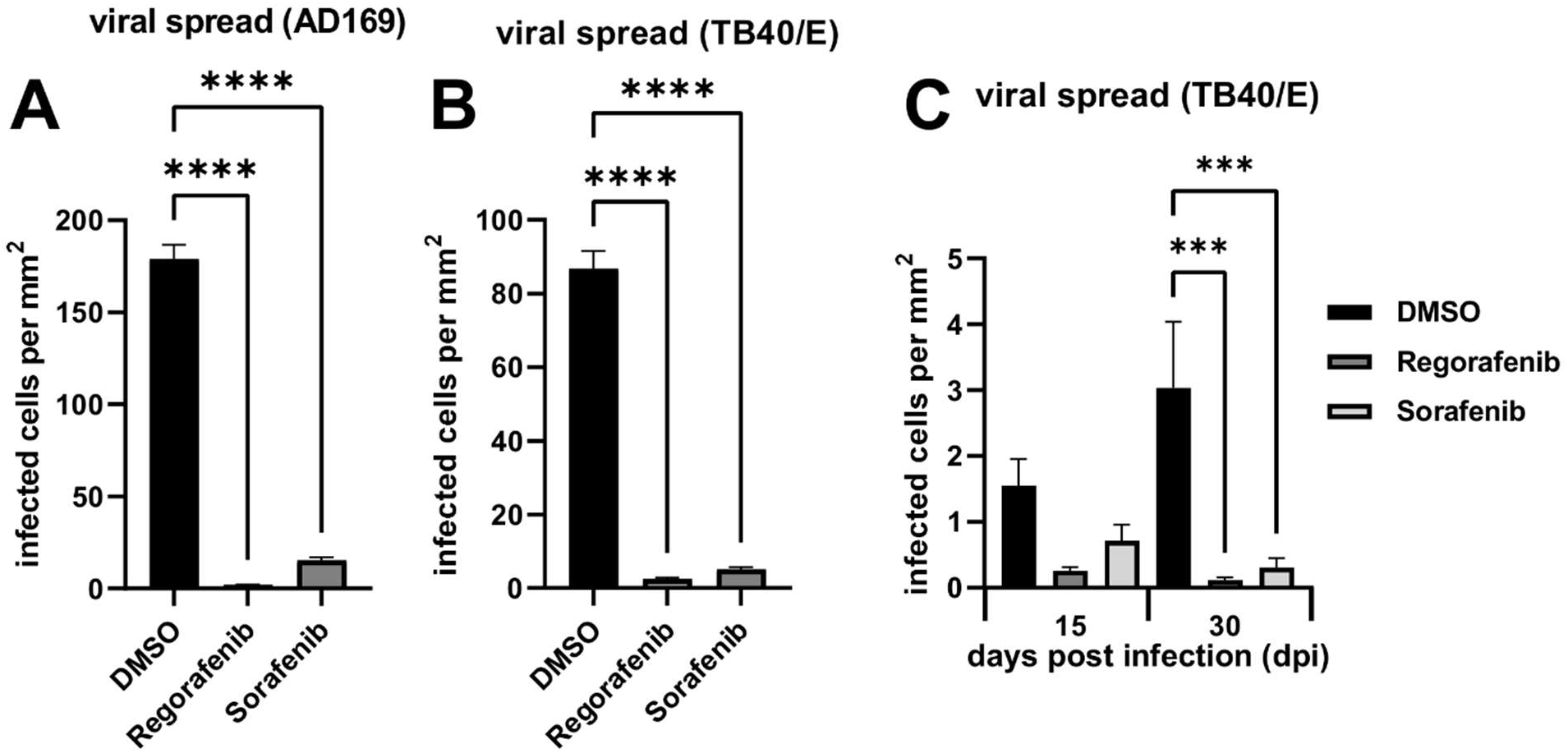
Pharmacological inhibition of Raf1 attenuates HCMV TB40/E cell-to-cell spread in fibroblasts and epithelial cells. (A-B) MRC5 fibroblasts or (C) ARPE19 epithelial cells were treated with 2.17 uM of Regorafenib or Sorafenib, or DMSO control at the time of infection with (A) AD169-GFP or (B-C) TB40/E-mCherry, at an MOI of 0.01. Viral spread was calculated by cells positive for (A) GFP or (B) mCherry on day 12 in MRC5hTs or (C) mCherry on days 15 and 30 (avg±SEM, n=48).

## Discussion

A variety of viruses rely on cellular growth factor signaling for many facets of viral infection including entry, the production of viral progeny, and the release of virions from the cell (20–24). These signaling molecules include various MAPK pathway activators and components, such as PDGFR, Ras, MEK, and ERK (23). Here, we show that Raf1 plays an important role in HCMV infection and is regulated by AMPK-mediated phosphorylation. Our data indicate that AMPK induces Raf1-specific phosphorylation during HCMV infection (**Figure 1**), which in turn enhances Raf1 binding to the 14-3-3 protein, an important co-factor for Raf1 activation (**Figure 2B-C**).

We find that pharmacological inhibition of Raf1 inhibits HCMV infection, reducing viral DNA accumulation and the production of viral progeny (**Figures 3 & 5**). Our results are consistent with a previous report that Sorafenib can inhibit HCMV infection in different cell types (36). Both Sorafenib and Regorafenib can inhibit multiple Raf kinases, including a-Raf, b-Raf, and Raf1, as well as VEGFR and c-Kit family members (**Table 1**) (49, 50). To address this issue, we targeted Raf1 more specifically via shRNA and CRISPR, and find that in both cases HCMV infection was reduced, albeit to a smaller extent than with pharmacological treatment (**Figures 6 & 7)**. These differences in the magnitude of viral inhibition indicate the possibility that other Raf family members may be able to support viral replication in the absence of Raf1, e.g., a-Raf or b-Raf, activities that would likely be inhibited by Sorafenib and Regorafenib.

Sorafenib and Regorafenib treatment inhibited HCMV replication in fibroblasts and epithelial cells (**Figures 3 & 8)**. Collectively, these results suggest the possibility of using these pharmacological Raf inhibitors to treat clinical HCMV infection. Regorafenib has FDA approval for the treatment of metastatic colorectal cancer, advanced gastrointestinal stromal tumors, and hepatocellular carcinoma. Sorafenib has FDA and European Commission approval for the treatment of hepatocellular, renal, and thyroid carcinomas. Both drugs are generally well tolerated in patients with mild side effects (51, 52). HCMV can be a significant pathological factor in cancer patients receiving immunosuppressive therapies (53–55). In this regard, the anti-viral activity of these compounds could therefore potentially help prevent HCMV-associated complications when they are used for cancer treatment.

We find that Raf1 phosphorylation during HCMV infection is complex. At early times post-infection, HCMV increases the levels of S338 phosphorylated Raf1 (**Figure 1**), consistent with its activation (39). By 48 hours post-infection the levels of Ser338 phosphorylation began to fall (**Figure 1**). The total levels of Raf1 also fell throughout infection (**Figure 1**). Further, Raf1 phosphorylation at Ser259 increased over the course of infection (**Figure 1**), which reportedly inhibits Raf1 activity (40). Collectively, these data suggest that the activity of Raf1 is reduced at the later times of infection. However, the functional relevance of potentially reduced Raf1 activity to infection is unclear. Of note, overexpression of Raf1 throughout infection did not negatively impact the production of viral progeny (**Figure 2**), suggesting that increased expression of Raf1 at late time points does not impact infection. However, we cannot rule out the possibility that Raf1 activity is still reduced at these late time points despite Raf1 over-expression, e.g., via inhibitory phosphorylation.

We assessed Raf1 phosphorylation using three phosphospecific antibodies that have been linked to Raf1 activation or inhibition. Notably, over fifty different Raf1 phosphorylations have been described in the literature (phosphosite.org), which highlights the difficulty of making broad conclusions based on data utilizing a few phosphospecific antibodies. Potentially capturing a broader representation of Raf1 post-translational modifications, HCMV infection induced dramatic changes to the isoelectric point of the majority of migrating Raf1 species, collapsing them and shifting them to a more acidic portion of the gradient (**Figure 1G**). These results are consistent with infection broadly inducing Raf1’s phosphorylation (**Figure 1G**). Further, treatment with Compound C substantially reversed this acidic shift, consistent with AMPK activity being important for Raf1 phosphorylation. Data using the Raf1 phospho-Ser621-specific antibody supported this possibility. HCMV infection induced Raf1 phosphorylation at Ser621, a site known to be phosphorylated by AMPK (47), and this phosphorylation was reversed with Compound C treatment (**Figure 1**). These data suggest that AMPK substantially modulates Raf1 phosphorylation during HCMV infection, yet many questions remain about how these AMPK-dependent modifications might functionally contribute to infection.

We found that AMPK-dependent Raf1-Ser621 phosphorylation increases as HCMV infection progresses (**Figure 1**). Studies suggest that Raf1 phosphorylation at Ser621 is required for its activation via binding to 14-3-3 binding (41, 56). Specifically, this phosphorylation facilitates ATP-binding upon 14-3-3 interaction (45). Consistent with this, HCMV infection induces Raf1 association with 14-3-3, which can be substantially reduced by a S621A mutation (**Figure 2**). Over-expression of Raf1-S621A did not impact HCMV infection, however firm conclusions about potential contributions of Raf1-Ser621 phosphorylation to infection are difficult given that wildtype Raf1 capable of Ser621 phosphorylation was present in these experiments.

AMPK has previously been found to be important for productive HCMV infection, in part through HCMV-induced metabolic remodeling, for example, inducing GLUT4 expression and activating glycolysis (18, 19). Here, our data suggest that AMPK might contribute to infection through modulation of Raf1 activity, a central MAPK signaling component that we find is important for infection. Collectively, our evidence suggests a link between HCMV-induced kinase pathways that are important for HCMV infection. Given HCMV’s reliance on these pathways, they may represent a therapeutically useful vulnerability, but questions remain as to how the cross-talk between these pathways might contribute to infection.

## Methods

### Cell and viral culture

Human 293T cells (ATC CCRL-3216), telomerase-expressing Human Foreskin Fibroblasts (HFF), telomerase-expressing MRC5 fibroblast cells, and HFF or MRC5 derivative cell lines were cultured in Dulbecco’s modified Eagle medium (DMEM; Invitrogen); ARPE19 retinal epithelial cells were cultured in Roswell Park Memorial Institute 1640 medium (RPMI; Invitrogen). MRC5 and HFF parental cell lines were generated via lentiviral transduction of telomerase, previously described here (37). DMEM and RPMI were supplemented with 10% fetal bovine serum (Atlanta Biologicals), 4.5 g/liter glucose, and 1% penicillin-streptomycin (Pen-Strep; Life Technologies) and cells maintained at 37°C in a 5% (vol/vol) CO2 atmosphere. Cell lines used for specific experiments are indicated in each figure legend.

Prior to HCMV infection, cells were grown to confluence and maintained in serum-free medium for 24 hours. Cells were mock- or HCMV-infected at the indicated multiplicity of infection (MOI), with HCMV viral strain AD169, AD169-GFP, or TB40/E-mCherry as indicated in the figure legends, for an adsorption time of 120 minutes. The AD169-GFP virus was generated as described here (57). The TB40/E-mCherry virus was a generous gift from Christine M. O’Connor and Eain Murphy, generated as described here (58). Afterward, the viral inoculum was removed, and cells were washed one time with Phosphate-Buffered Saline (PBS; Invitrogen) and subsequently cultured in serum-free DMEM or RPMI for the duration of infection unless otherwise indicated. For viral spread assays, cells were grown in a 384-well plate; after adsorption, the medium was not aspirated and cells were not washed with PBS, but wells were brought up to the final well volume of 7 µl for the remainder of the experiment. All drug compounds, when indicated, were added at the time of viral adsorption and again with replacement media after viral inoculum was removed for the remainder of the experiment. All viral stocks were propagated in MRC5 cells and maintained in serum-free DMEM with Pen-Strep present. Viral stock titers were calculated by plaque assay.

### Flag-Raf1 plasmid cloning

The Flag-Raf1-WT and Flag-Raf1-S621A constructs were cloned via Gibson assembly into the pLenti-puro vector. The Raf1 sequence was taken from the pDONR223-RAF1 construct ordered from Addgene, a gift from William Hahn & David Root (Addgene plasmid # 23832; http://n2t.net/addgene:23832; RRID:Addgene_23832). The FLAG tag was inserted with a Gibson assembly primer. The S621A mutation was introduced via Gibson assembly into the wild-type (WT) construct. Assembled constructs were heat shocked into Stbl3 bacterial cells for plasmid propagation and storage.

### shRNA knockdown

For shRNA knockdown experiments, pLKO.1-based Mission shRNA constructs targeting Raf1 were obtained from the Sigma-Aldrich Mission shRNA library (Sigma/Broad Institute; clone number TRCN_1065 (shR1-1) or TRCN_1068 (shR1-5)). Mission pLKO.1-puro scrambled non-targeting empty vector (EV) control shRNA (EV shRNA; SHC002) was used as a control in all shRNA experiments. The effectiveness of each shRNA construct to knockdown Raf1 was assessed by qPCR.

### Lentiviral transductions and cell line generation

For each Raf1 lentiviral construct, three 10 cm dishes were seeded with 293T cells at 2×10^6^ cells per dish; similarly, one 10 cm dish was seeded for each shRNA construct. After 24 hours, each dish of cells was transfected with 2.6 μg lentiviral transfer or shRNA construct plasmid, 2.4 μg PAX2 packaging plasmid, and 0.25 μg VSV-G envelope plasmid using Fugene 6 (Promega) transfection reagent. After an additional 24 hours, the medium was removed and replaced with 4 ml of fresh medium for 24 hours. The following day, the supernatant was filtered through a 0.45 μm syringe filter to remove any debris and dead cells. Filtered lentiviral supernatant was introduced to MRC5 cells for FLAG-Raf1 experiments or HFF cells for shRNA experiments in the presence of 5 μg/ml Polybrene (Millipore Sigma). Three hours later, the lentivirus on FLAG cells was removed, and freshly filtered lentivirus from a second dish of 293T cells was added without polybrene. After 3 more hours, lentivirus was once again removed and replaced with freshly filtered lentivirus from the final dish of 293T cells without polybrene and incubated overnight. Plates transduced with the shRNA lentiviral constructs were left overnight. The following day, lentivirus was replaced with 10 ml of fresh DMEM containing FBS and Pen-Strep. 72 hours post-transduction, cells were selected with 1 μg/ml puromycin in normal growth conditions. Selection efficiency/efficacy was monitored by cell death of mock-transduced parental cells. Once mock cells were dead, shRNA-transduced cells were immediately used for experiments. Flag-Raf1 cells were used immediately, and some cells were frozen down for later use. If thawed, the expression of Flag-Raf1 was verified by Western blot before experiments were carried out.

### CRISPR knockout cell line generation

CRISPR synthetic single guide RNAs (sgRNA) and SpCas9 2NLS Nuclease were ordered from Synthego Corporation. Guide sequences are listed in **Table 2**. For each transfection, 30 pmol of the sgRNA and 10 pmol of Cas9 were introduced to 2X10^5^ MRC5 or HFF cells via the Neon transfection system (Invitrogen) and the Neon transfection system 10 ul kit according to the manufacturer’s protocol. Neon electroporation parameters for the MRC5 fibroblasts were as follows: Voltage - 1100, pulse width – 30 ms, 1 pulse; and for the HFF cells: Voltage - 1650, pulse width – 10 ms, 3 pulses. Parental cells were shocked via Neon using the same parameters, with equivalent volumes of buffer and cell suspension. After electroporation, cells were immediately plated into prewarmed DMEM with FBS and no antibiotics overnight in a 12-well plate. Cells were propagated as normal once they became confluent. Primers used to sequence genomic DNA for ICE analysis (Synthego Corporation) to determine the KO score of each polyclonal cell line are listed in **Table 3**.

**Table 2.**
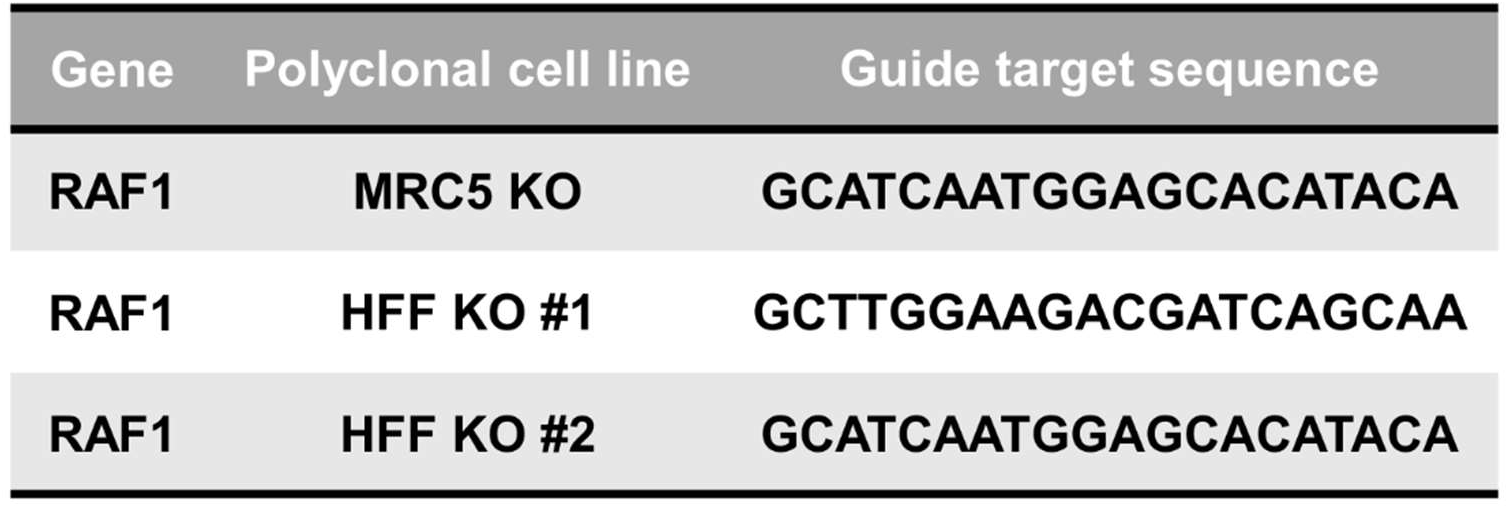
CRISPR guide sequences.

**Table 3.**
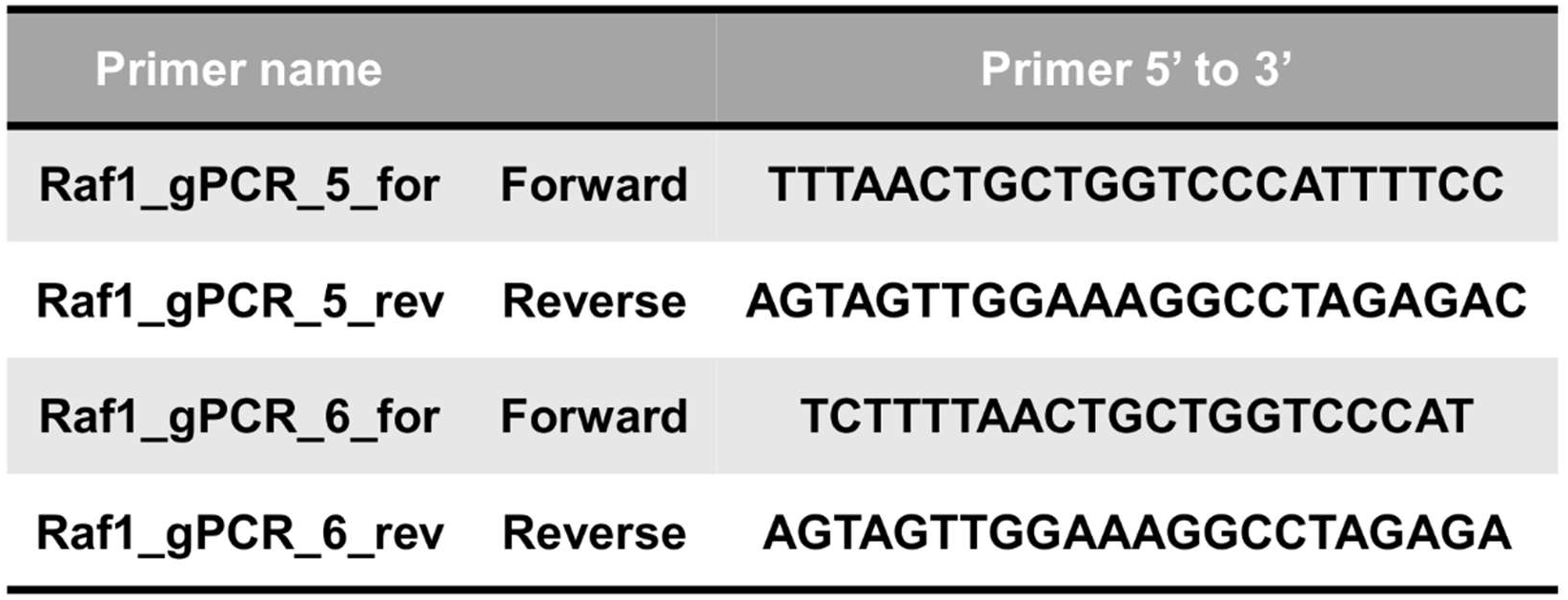
Primers used for genomic PCRs of CRISPR polyclonal knockout cell lines.

### Drugs and compounds

All drugs were suspended in vehicle control, dimethyl sulfoxide (DMSO; Corning). Compound C (CC; Millipore Sigma or MedChemExpress) was purchased in solution at a concentration of 10 mM in dimethyl sulfoxide (DMSO), stored at -80 or - 20°C for less than 2 months prior to use, and used at a concentration of 5 μM for all experiments. Regorafenib and Sorafenib (MedChemExpress) were suspended in DMSO and used at a concentration of 2.17 µM unless otherwise indicated. Hoechst 33342 (Invitrogen) was purchased in solution at a concentration of 10 mg/ml and was diluted 1:2,000 in PBS for use. Puromycin (Millipore Sigma) was purchased as a powder, suspended to 10 mg/ml in water, and used at a final concentration of 1 μg/ml for all lentiviral transduction selections.

### Immunoblotting

Cells were harvested by washing once with cold PBS, then scraping on ice in 1X RIPA (10X RIPA: 0.5 M Tris-HCl pH 7.4, 1.5 M NaCl, 10 mM EDTA, 2.5% deoxycholic acid, 10% TritonX-100) or cold lysis buffer (20 mM Tris-HCl (pH 7.5), 100 mM NaCl, 1 mM MgCl2, 1% IGEPAL CA-630, 1 mM EDTA) containing a protease inhibitor tablet (Pierce; A32955) and PhosSTOP tablet (Sigma) per 10 ml of buffer. RIPA was used for all Western blots, apart from the Raf1 overexpression experiments, where lysis buffer was used. Samples were incubated in a tube on ice for 20 minutes, sonicated, then the insoluble fraction was pelleted and discarded. Soluble proteins were quantified via Bradford assay (BioRad) and samples were normalized to total protein concentration and added 3-parts protein to 1-part 4X disruption buffer (200 mM Tris-HCl (pH 7.0), 11% sucrose, 20% beta-mercaptoethanol, 8% SDS), then to boiled for 5 minutes. Samples were spun down prior to running on a gel.

For flag pulldown experiments, protein lysates were added to 50 ul of prewashed ANTI-FLAG M2 Affinity Gel beads (Sigma) and incubated at 4°C overnight with gentle rotation. Pulldowns were washed 4 times with modified lysis buffer (IGEPAL CA-630 reduced to 0.1%). Then 1X disruption buffer was added to washed beads and boiled for 5 minutes. Samples were spun down and the disruption buffer containing immunoprecipitated proteins was transferred to a new tube.

Proteins were separated on a 10% SDS-PAGE gel in Tris-glycine running buffer. Samples were transferred to a nitrocellulose membrane in Tris-glycine transfer buffer and stained with Ponceau S to visualize proteins and ensure equal loading. Blots were blocked with 5% dried non-fat milk in Tris-buffered saline-Tween 20 (TBST), followed by primary and secondary incubations in 5% Bovine Serum Albumin (BSA) in TBST (Flag primary was diluted in 2.5% milk in TBST). Antibodies were used at concentrations recommended by the manufacturer and are listed in **Table 4**. Blots were developed with enhanced chemiluminescence (ECL, Bio-Rad) and imaged using a Molecular Imager Gel Doc XR+ system (Bio-Rad). Western blot densitometry was measured using Bio-Rad Image Lab Software taking the total band intensity.

**Table 4.**
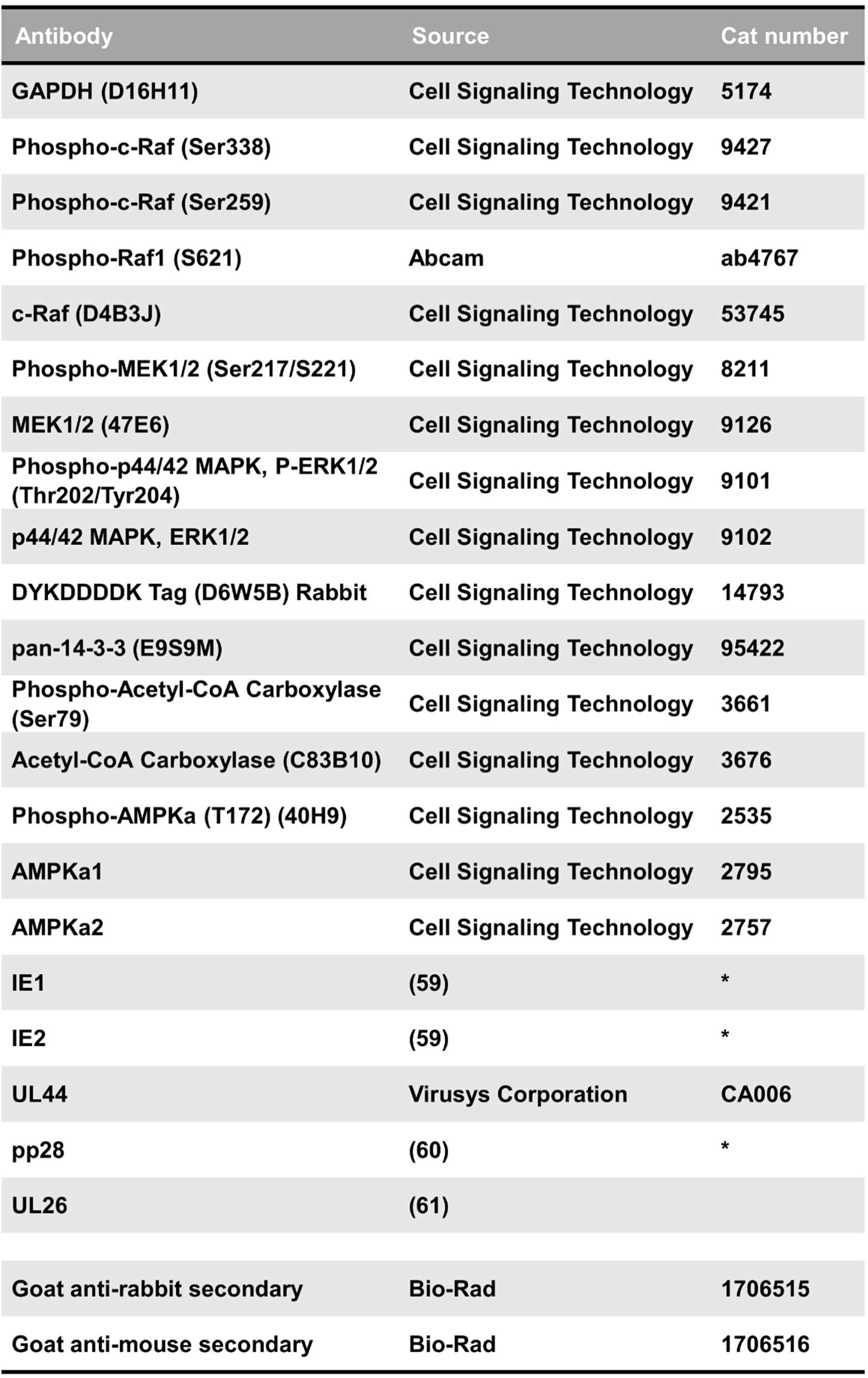
List of antibodies used. *Generous gift from Thomas Shenk.

### Two-dimensional (2-D) gel electrophoresis

Cells were treated with Compound C and infected in 15 cm dishes, then 2-D gel samples were collected by scraping cells in 250 ul of 2-D rehydration buffer (4% CHAPS, 8 M urea, 0.5% IPG Buffer (pH 3.0-10.0; GE Healthcare), 0.002% Bromophenol Blue, 40 mM DTT (added fresh before extraction)). Samples were spun to pellet insoluble fraction and loaded on a 13-cm Immobline Drystrip isoelectric-focusing gel (pH 3.0-10.0; GE Healthcare), and proteins were separated in the first dimension using the IPGphor isoelectric-focusing system (Amersham Pharmacia Biotech) following manufacturer’s protocol. Then gel was equilibrated for 15 minutes in SDS equilibration buffer (6 M urea, 75 mM Tris-HCl (pH 8.8), 30% Glycerol, 2% SDS, 0.002% Bromophenol Blue), and proteins were run in the second dimension by SDS-based electrophoresis on a 10% polyacrylamide gel. Finally, samples were treated as a typical western blot, by transferring the proteins to a nitrocellulose membrane, blocking with milk, and immunoblotting for Raf1 as described above.

### Cell-to-cell spread assay and nuclear counts

For the drug titrations performed in **Figure 3**, cells were seeded using the Multidrop Combi Reagent Dispenser (Combi; Thermo Scientific) into a 384-well plate and grown to confluence. The medium was then aspirated and replaced with serum-free medium for 24 hours. Then cells were treated with Regorafenib and Sorafenib using the D300 digital dispenser (D300; Hewlett-Packard Company, L.P) with accompanying software to titrate into the low nM range. Immediately afterwards AD169-GFP virus was dispensed using the Combi. After 2 hours, all wells were brought up to the final culture volume with fresh medium and the drug for the remainder of the experiment.

For cell spread assays in **Figure 7D-E**, and **Figure 8**, indicated cell lines were seeded into 384-well plates and grown to confluence. The medium was then aspirated and replaced with serum-free medium for 24 hours. Then cells were treated with 2.17 uM drug or volume equivalent of DMSO control (using the VOYAGER automated digital pipette; Integra Biosciences) when indicated (**Figure 8**) and immediately infected with AD169-GFP or TB40/E-mCherry as indicated using the Voyager pipette for 2 hours. All wells were brought up to the final volume with drug or DMSO control for the remainder of the experiment.

GFP or mCherry-positive cells were imaged on the Cytation 5 imaging reader and masked using the accompanying Gen5 software (BioTek Instruments, Inc.). The final cell-to-cell spread was calculated as GFP or mCherry-positive cells per area of the entire well. Cell counts in **Figure 4** were acquired with Hoechst 33342 (Invitrogen) staining of nuclei, followed by imaging on the Cytation 5 imaging reader. Nuclei were counted using the Gen5 software.

### Real-time quantitative PCR (qPCR)

RNA was extracted using Trizol reagent (Invitrogen) according to the manufacturer’s protocol, RNA was treated with DNase I (Invitrogen), and used to synthesize cDNA using random hexamer primers (Invitrogen) and Superscript II reverse transcriptase (Invitrogen). Quantitative PCR (qPCR) was carried out using Fast SYBR Green Master Mix (Applied Biosystems), using a 7500 fast real-time PCR system (Applied Biosystems), and StepOne real-time PCR software (Applied Biosystems). Primers used for each reaction are listed in **Table 5**. Raf1 primers were obtained from PrimerBank, ID:189458830c2 (59). Relative RNA levels for Raf1 were measured and normalized to GAPDH levels and then indicated shRNA-NT samples, using the 2-ΔΔCT method.

**Table 5.**
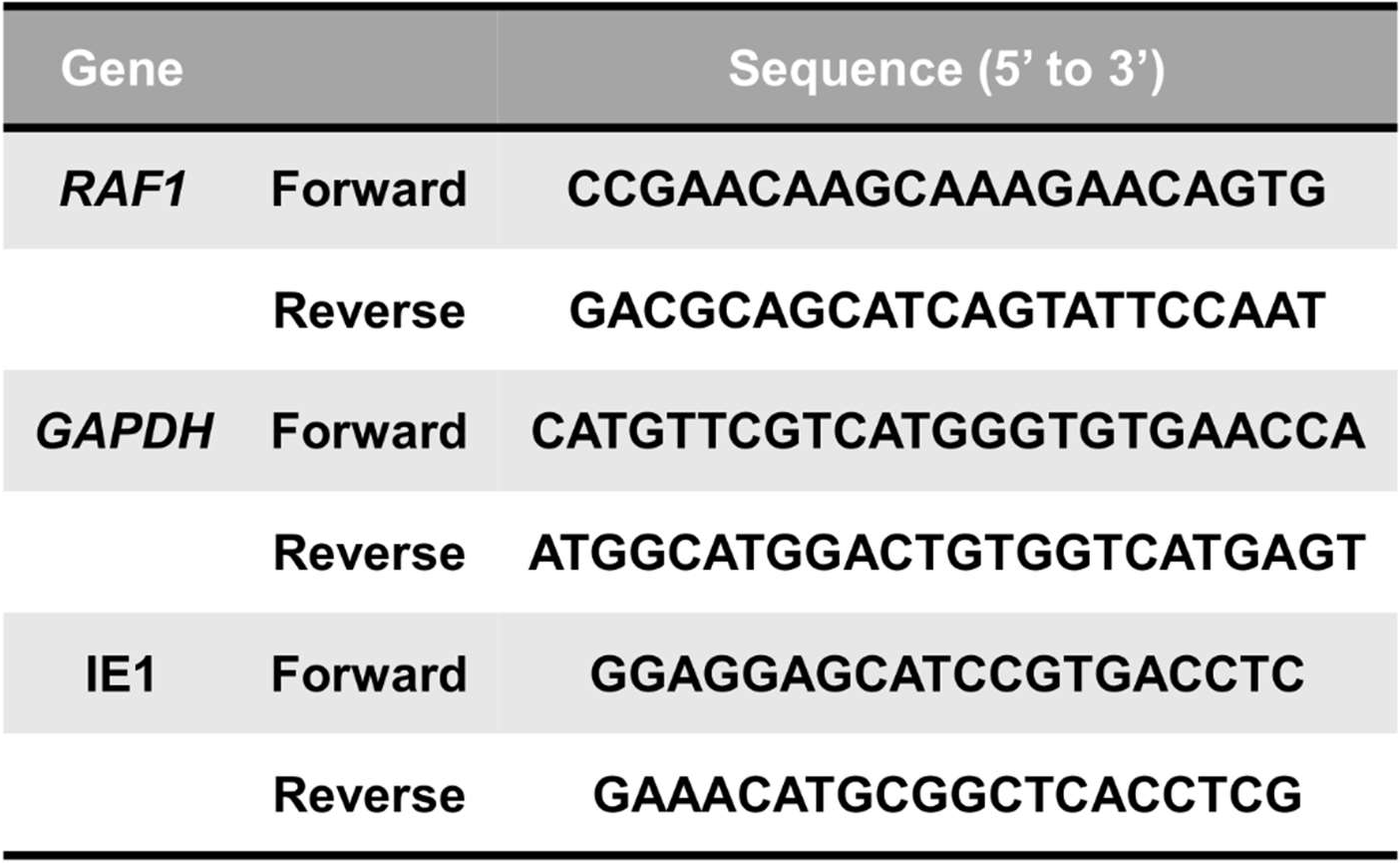
List of qPCR primer sequences.

### Statistical analysis

Statistical analysis was carried out using GraphPad Prism 9 statistical software, sample number (n) ± standard deviation (STD) or standard error mean (SEM) are plotted as indicated in figure legends. A one-way ANOVA or Student’s t-test was performed as indicated in the figure legends. P<0.05 was considered statistically significant, where * is P<0.05, ** is P<0.01, *** is P<0.001, and “ns” indicates not significant. Area under the curve (AUC) was calculated using GraphPad Prism 9 software. IC50 values reported in **Figure 3** were calculated using GraphPad Prism 9 using the nonlinear regession (curve fit), [Inhibitior] vs. response -- variable slope (four parameters).

## Acknowledgments

The work was supported by NIH grants AI127370 and AI50698 to J.M. D.M.D. was supported by a postdoctoral fellowship (133137-PF-19-038-01-MPC) from the American Cancer Society. The schematic in **Figure 1A** was created in biorender (biorender.com).

D.M.D. performed the experiments, prepared the figures, performed statistical analysis, and formatted the manuscript and figures for submission. L.J.P. cloned the pLenti-Flag-Raf1-WT construct. The authors would like to thank Katie Tomberlin for validating the qPCR primers used to quantify the shRNA knockdown during her graduate school rotation in J.C.M.’s laboratory. The authors also thank Xenia L. Schafer for proofreading and editing this manuscript. D.M.D. and J.C.M. contributed to the conceptualization of this project, designed the experiments, and wrote the manuscript. All authors, D.M.D., L.J.P., and J.C.M., contributed to the editing of the manuscript.

